# Variation in mechanical and structural properties of enamel in primate molars

**DOI:** 10.1101/2021.07.05.451217

**Authors:** Ian Towle, Thomas Loho, Amira Salem, Carolina Loch

## Abstract

Mechanical properties of enamel are known to vary across molar crowns in some primates, but the association of this variation with phylogeny, structural properties and tribological behaviour is not well understood. In this study, 20 molars from a range of primate taxa (n=15) were studied using nanoindentation, micro-CT scanning, and SEM imaging. After micro-CT scanning, teeth were sectioned in the lingual-buccal plane through the mesial cusps. Five positions (buccal lateral, buccal cuspal, occlusal middle, lingual cuspal, lingual lateral) were studied in three locations (inner, middle, outer enamel regions). The results show middle enamel had the highest hardness and elastic modulus values in all positions. ‘Non-functional’ molar sides (lingual in lower molars and buccal in upper molars) had higher hardness values than their ‘functional’ counterparts. Increase in prism size was associated with a decrease in hardness in some tooth positions, and mineral density showed a significant relationship with elastic modulus values. Variation in enamel structure variation (e.g., enamel Schmelzmuster, Hunter-Schreger band thickness), may also be crucial in explaining variation in mechanical properties, with decussation zones associated with higher mechanical properties values. Primate enamel is not a homogeneous material, with variation in mechanical and structural properties across the crown likely associated with functional differences and variation in force distribution. Overall structural and mechanical patterns were similar in the primate species studied despite substantial differences in diet, suggesting these properties are potentially evolutionary conserved.

## Introduction

Enamel forms the outer layer of molar crowns in primates and is often characterised by its durability, being the hardest biological material in the body (He and Swain, 2008; Jeng et al., 2011). Enamel consists of both inorganic and organic components, although organic components make up less than 5% of the enamel volume (Cuy et al., 2002). The organisation and structure of enamel is complex and can be studied at a variety of scales, from individual crystals, through prisms, HSB and the overall properties of the enamel in the crown. Previous research on enamel evolution in primates has typically investigated the distribution of enamel on the crown and its overall thickness, rather than structural, mechanical and compositional variation on a smaller scale.

It is well known that enamel is not evenly distributed in primate molars, being substantially thicker on the lingual side of upper molars, and buccal side of lower molars (Kay, 1975; Molnar and Ward, 1977; Macho and Berner, 1993; Schwartz, 2000). The cusps on the side with thicker enamel are often termed ‘functional’ cusps, i.e. the buccal of lower molars and the lingual of upper molars, with the opposite surfaces termed ‘non-functional’ cusps (Khera et al., 1990; Schwartz, 2000). The extent to these differences varies among primate groups. In colobines, the enamel thickness is more homogeneous between buccal and lingual cusps than in other primates (Ulhaas et al., 1999). Previous studies have suggested that thicker enamel on functional cusps may relate to protection against fracture (Grine, 2005). However, other studies have suggested that thicker enamel in these cusps may relate to managing shearing forces, with nonfunctional molar cusps dealing with crushing forces (Schwartz, 2000). In support of the later hypothesis, functional cusps seem to show more rapid wear (Macho and Berner, 1993; Schwartz, 2000; Kono, 2002), and non-functional cusps more chipping (Cavel et al., 1985; Eakle et al., 1986; Towle et al., 2021). Despite this, functional cusps are usually associated with higher stress loads than their nonfunctional counterparts (Kay, 1975; Lucas et al., 2008; Thiery et al., 2017).

Little is known about differences in mechanical and structural properties of enamel between crown locations, including between functional and non-functional cusps. Typically, in comparative studies on the biomechanics of enamel, it is often considered a homogeneous material. Only a small number of studies have looked at how mechanical properties (i.e., hardness and elastic modulus) vary across primate molar crowns in relation to changes in structural and compositional factors (e.g., prism and interprismatic matrix organisation, *Schmelzmuster*, mineral composition and mineral concentration). These studies have made it clear enamel is not homogeneous, with substantial variations identified between locations in the same tooth crown (Cuy et al., 2002; Park et al., 2008a; Lee et al., 2010; Darnell et al., 2010; Campbell et al., 2012; Constantino et al., 2012).

In these mechanical property studies, hardness and elastic modulus values were typically collected using a nanoindenter, with hundreds or even thousands of individual indents recorded across the crown. One of the main variations in mechanical properties found within primate enamel has been hardness and elastic modulus increasing from inner enamel (near the CEJ) to outer enamel, which appears particularly consistent in humans (e.g., Cuy et al., 2002; Park et al., 2008b). However, other studies have suggested this may not be the case for certain primate groups (Campbell et al., 2012; Constantino et al., 2012). Other changes in mechanical properties across tooth crowns have also been observed in these studies, including variation between buccal/lingual and lateral/cuspal positions (Cuy et al., 2002; Darnell et al., 2010). Factors that may influence variation in enamel mechanical properties have been investigated mainly in humans, with factors such as mineralization, prism orientation, enamel composition, and crystal arrangement/orientation being considered (e.g., Cuy et al., 2002; Angker et al., 2004; Shimizu and Macho, 2008; Xie et al., 2009; Jeng et al., 2011; Stifler et al., 2021).

Overall mean mechanical property values often differ between species (e.g., Kaiser et al., 2018), although other studies have shown there is little variation in mechanical properties between primate taxa (Constantino et al., 2012). Other properties, such as occlusal morphology and/or enamel thickness, may show more phylogenetic variation related to changes in diet (Lee et al., 2010; Constantino et al., 2012). It is likely that variation in mechanical properties across individual tooth crowns may relate to regional differences in stress distribution (Cuy et al., 2002). This is supported by research showing that differences in enamel properties (e.g., microstructure, composition, hierarchical structure) in different crown locations will affect the likelihood of tooth wear and fracture (Zheng et al., 2003; He and Swain, 2008; Roy and Basu, 2008; Jeng et al., 2011; Barani et al., 2012; Constantino et al., 2012). There are also clear differences in enamel structure and composition between mammals with varied feeding behaviours (e.g., Xiao et al., 2019).

This study aims to expand on previous research in enamel mechanical properties and variation across tooth crowns in a wide range of primates. To achieve this, different regions of the enamel were investigated (outer, inner and middle), in five locations (two buccal and lingual locations and one occlusal). The aim was to test if changes in mechanical properties have a phylogenetic signal, and if differences in properties in functional and non-functional sides might relate to wear and fracture patterns (e.g., Towle et al., 2021). Factors such as prism width, enamel thickness, mineral concentration, total effective density and enamel *Schmelzmuster* were considered. Based on previous studies and assumptions based on the functional adaptations of teeth, this study tested three hypothesis: 1) hardness and elastic modulus values will increase with increase in enamel thickness (e.g., ‘functional’ cusps will have higher values); 2) there will be an increase in hardness and mineral density from inner to outer enamel and this will be associated with increase in prism width; 3) cuspal enamel will show higher values for enamel properties than lateral enamel, but overall tooth averages will be similar among studied taxa.

## Materials and methods

### Samples

A range of Catarrhini (n=19) and one Platyrrhini species were analysed. Most specimens are assigned to species level; however, some teeth were assigned to genus to avoid misidentification. The 20 teeth analysed here are composed of 16 lower molars and four upper molars. Most teeth studied were second (n=14) and first molars (n=6). Unworn or moderately worn teeth were selected when possible. Analysed teeth also did not show visible pathology, enamel defects or macroscopically visible post-mortem damage. Specimens belong to collections in New Zealand (Otago Museum; Museum of Natural Mystery; Auckland Museum; Faculty of Dentistry, University of Otago), and the Primate Research Institute, Kyoto University, Japan. Information on each specimen is presented in Supplementary Table 1. Prior to testing, all teeth were stored dry, except for the two human teeth which were stored in formaldehyde. These human teeth were extracted as part of planned clinical treatment and were obtained with patient consent.

Each tooth was first micro-CT scanned, followed by sectioning for nanoindentation and SEM analyses. For each protocol (micro-CT scans, nanoindentation and SEM) readings were taken for 15 locations within the buccal-lingual plane of the mesial cusps (described below). These locations are within the inner, middle and outer enamel in five positions: lingual lateral; lingual cuspal; middle occlusal; buccal cuspal; buccal lateral (Figure 1). Areas with cracks, wear and other artefacts were avoided. The positions were standardised as follows, with buccal and lateral referring to the two sides of the mesial cusps: 1) lateral: within the lower half of the crown, but before enamel begins to taper towards the cementum-enamel junction; 2) cuspal: approximately a straight line from the maximum height of the cuspal dentine to the outer enamel (if cusps were slightly worn, a cervical point was chosen); 3) approximately the middle of the occlusal surface (i.e., halfway between the two cusps in the central fissure).

**Figure 1.**
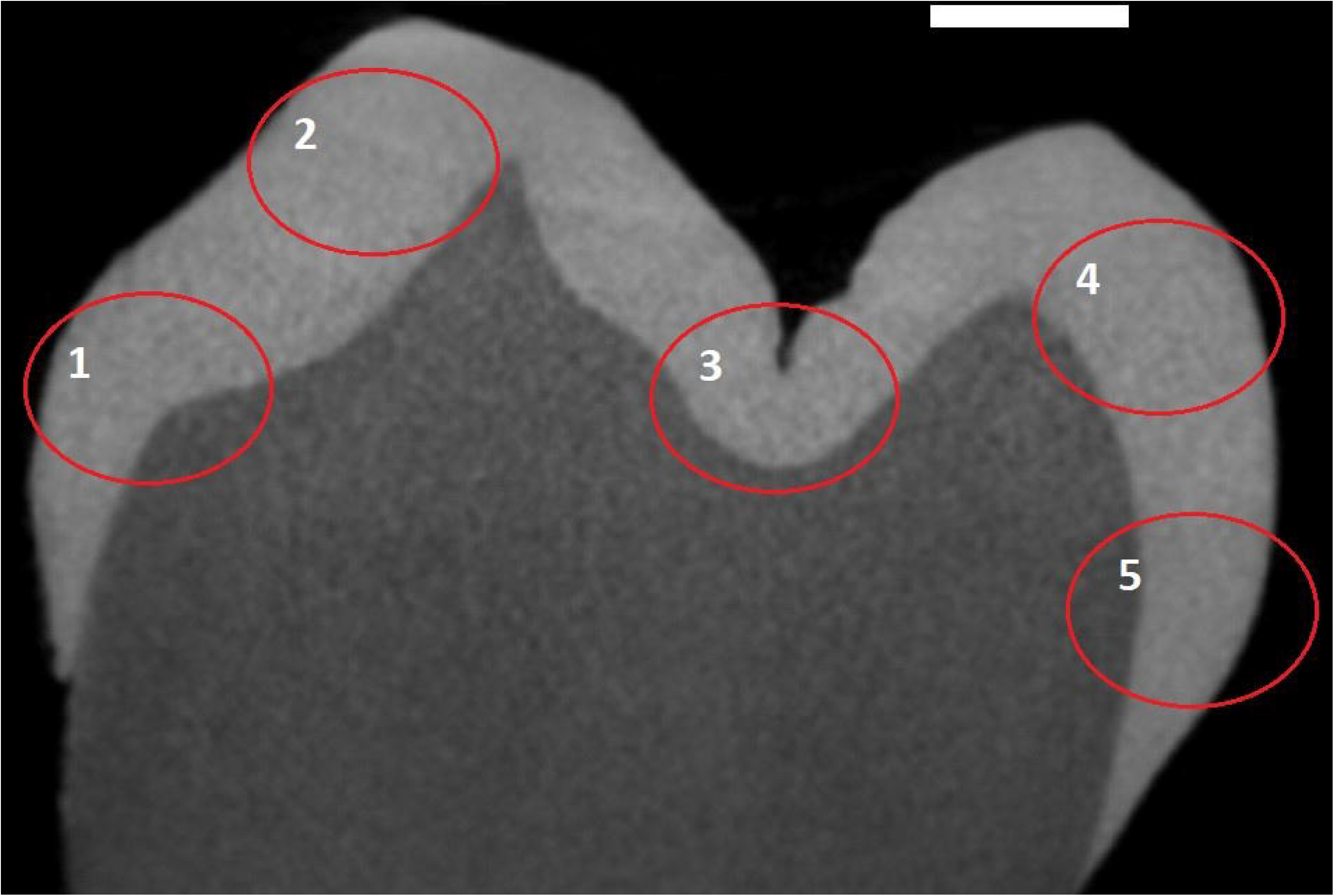
Sampling locations and the five positions studied on a lower human molar. Slice from buccal (left) to lingual (right), through the middle of the mesial cusps. Positions: 1) buccal lateral; 2) buccal cuspal; 3) middle occlusal; 4) lingual cuspal; 5) lingual lateral. Scale bar: 1mm.

### Micro-CT scanning

Teeth were scanned with a Skyscan 1172 Micro-CT desktop system (Skyscan, Kontich, Belgium). X-rays were generated at 100 kV, 100 μA and 10W. 0.5 mm thick aluminium and copper filters were placed in the beam path. The resolution was set at 15 μm pixel size and rotation was set to 0.2-degree for all teeth. Images were reconstructed using the Skyscan NRecon software (NRecon, version 1.4.4, Skyscan) in order to create cross-sectional slices of the specimens. Reconstruction settings were standardized as follows: smoothing 5; ring artefact correction 20; beam hardening correction 80%. Resin-hydroxyapatite phantoms were used to calibrate the grayscale and mineral density of enamel in each specimen (following Schwass et al. 2009). Grayscale values were recorded on each of three density standards using 3mm oval Regions of Interest (ROI’s) on ImageJ, with three positions recorded on every 10 slices. Average grayscale values for each phantom along with the known Mineral Concentration (MC) and Total Effective Density (TED) were then used to produce a linear regression for each specimen with the resulting equation used to calculate MC and TED values for each enamel location. For MC, the R^2^ was consistently high among samples for the three phantoms (R2= 0.99 ± 0.007). A fourth point was added to the TED linear regression, using an assumed point at which MC and TED converge at 3 g/cm3 (following Schwass et al. 2009), which increased the R^2^ and lowered the standard deviation (three points: R2= 0.95 ± 0.016; four points: R2= 0.98 ± 0.001). This fourth point was calculated for each sample using the MC linear regression equation to calculate the grayscale value at 3g/cm3.

Scans were calibrated for size (mm) and density using the ‘set scale’ and ‘calibrate’ functions in ImageJ. After calibration with the phantoms, MC and TED was calculated for each enamel location, using an oval ROI with a diameter of 0.15mm. 15 locations were sampled in each slice, which includes outer, middle and inner readings for the five enamel locations across the crown (lingual lateral; lingual cuspal; middle occlusal; buccal cuspal; buccal lateral). Moving distally from the mesial interproximal surface of the tooth, data collection started at the slice in which all five positions could be recorded. Moving distally through the tooth, data was collected every 10 slices until the end of the mesial cusps. This meant 15 to 30 slices were used per specimen, depending on the size of the tooth. Areas with post-mortem damage (e.g., cracks) or scanning/processing artefacts (e.g. ring artefacts) were avoided.

### Sample preparation

For the destructive analysis, preparation protocols followed Loch et al. (2013a/b), and are summarised below. Before embedding in resin, tooth surfaces were cleaned with ethanol. Each tooth was placed in a silicone mould, and was oriented to allow sectioning through the buccal-lingual plane of the mesial cusps. Teeth were embedded in epoxy resin (Epofix™ Cold-Setting Embedding Resin, Struers, Copenhagen, Denmark). After setting at room temperature for 24 h, teeth were longitudinally sectioned through the buccal–lingual plane of the mesial cusp tips using a MOD13 diamond blade (Struers, Copenhagen, Denmark) under water irrigation in a high-speed saw (Accutom-50, Struers, Copenhagen, Denmark). Two tooth segments were produced (distal and mesial), with the mesial side being used for the nanoindentation testing and the distal for SEM imaging. Distal and mesial sections were cleaned with ethanol and polished on a TegraPol-21 polisher (Struers, Copenhagen, Denmark) with 1200 grit silicon carbide paper (Struers, Copenhagen, Denmark) and ultrasonic cleaned for 3 minutes. Specimens prepared for SEM imaging (distal segment) were further polished with 2400 and 4000 grit silicon carbide paper, and ultrasonic cleaned for 3 min in water. After polishing, specimens were ultrasonically cleaned in ethanol for 3 min before etching with 2M phosphoric acid for 5-10 sec. Specimens were sonicated again in ethanol for 3 min. Specimens prepared for nanoindentation testing (mesial segment) were further polished with 1μm diamond suspension (DP Suspension P, Struers, Copenhagen, Denmark). Final ultrasonic cleaning was done in ethanol for 3 min.

### Nanoindentation testing

After polishing, the mesial half of each specimen was mounted on a steel base using thermoplastic cyanoacrylate glue. The mounting base contained a strong magnet to prevent any lateral movement during nanoindentation. A 20x magnification objective lens optical microscope coupled with the nanoindenter allowed precise positioning for each indent array. Tests were performed using a Hysitron TI 950 Triboindenter (Bruker, Minnesota, USA) equipped with a three-plate capacitive transducer capable of applying 10 mN of force with a 1 nN resolution, and 5 μm displacement with a 0.04 nm resolution. Each indent was performed using a three-sided pyramidal Berkovich tip (Bruker, Minnesota, USA) that was calibrated on a fused quartz standard. The load function for each indent was a load-controlled quasi-static trapezoid load function with 5 sec loading time, 2 sec holding time, and 5 sec unloading time with a maximum load of 10 mN. Indent arrays were standardized across samples, with 3 rows containing 15 indents in each of the five positions, giving a total of 225 indents per specimen. Typically, the last 2-4 columns indented into dentine, and were removed before analysis, leaving approximately 165-180 enamel indents per tooth. For each position, the first row of indents was undertaken slightly in (≈ 10 μm) from the outer enamel surface. Due to variation in enamel thickness between positions and among species, the distance between indents was varied to obtain at least 165 indents within the enamel for each sample. A minimum distance of 15 μm was set to avoid the influence of residual stresses from adjacent previous indents. Indents were later proportionally divided as belonging to the inner, mid or outer region, meaning approximately 11-13 indents for each position from which a mean was calculated. The reduced modulus (Er) and hardness (H) values were calculated using the TriboScan 9 software (Bruker, Minnesota, USA) based on the Oliver–Pharr method (Oliver and Pharr, 1992). Calculation of elastic modulus (E) from reduced modulus values assuming a Poisson’s ratio of 0.30 for enamel showed that the elastic modulus values were similar to the reduced modulus values (mostly less than 1.0 % difference). Thus, the reduced modulus was reported as the elastic modulus in this study. All nanoindentation tests commenced after thermal equilibrium was achieved in the instrument and tests were performed under standard laboratory conditions (approximately 21°C and 56% relative humidity).

### SEM imaging

Polished samples were coated with gold palladium prior to SEM analysis. A JEOL JSM-6700F Field Emission SEM (JEOL Ltd., Tokyo, Japan) was used operating at 5kV and 10 μA. Images were taken in the same testing positions as above. Prism size was recorded following Campbell et al. (2012), within each of the five locations, in areas of prismatic enamel in which prisms are exposed roughly perpendicular to the section were selected, to reduce the effects of section angle on prism width. Ten measurements were taken for inner, middle and outer enamel within each of the five positions (total of 150 readings per tooth). For prism width measurements, the magnification was set at 2,000x. For assessing the *Schmelzmuster* of enamel, the magnification ranged from 300 to 3,000x. An overview image of each position was also taken, using this magnification range, to assess the presence of Hunter-Schreger Bands and other features.

### Comparisons and analysis

The five positions were compared to assess the association between prism width, *Schmelzmuster*, MC/TED and enamel thickness with mechanical properties values (hardness and elastic modulus). Specimen averages for inner, middle and outer enamel were also compared for all variables. Additionally, comparisons were made based on *Schmelzmuster* differences between species (e.g., variation in Hunter-Schreger band extent across the enamel thickness). Comparisons were made between cuspal and lateral positions, and between buccal and lingual positions. For comparisons using different methods (e.g., TED Vs. hardness) mean values for each position were compared using a Pearson Correlation test with significance set at 0.05. A two-tailed T-test was used to compare the same variable (e.g., elastic modulus) in different crown positions (e.g., buccal Vs. lingual cusps). Results were also compared to chipping and wear variation between these same locations using published data (Towle et al., 2021; Towle and Loch, 2021), to explore how variation in mechanical properties and structure may relate to different types of hard tissue loss. Unless stated otherwise, the data used for statistical testing were the average values for each of the three sites (inner, middle, outer) for the five positions (i.e., 15 values per sample). The mean value for each variable, for each sample, was also tested against body mass, to ascertain if increase in body size may be related to changes in tooth properties. Average body mass data for each taxon followed Galán-Acedo et al. (2019), with average body mass values for genera used for samples in which species level identification was not possible.

## Results

The overall values for mineral density and mechanical properties variables show similar averages among samples, consistent with their close phylogenetic relationships (Table 1). Average values for hardness ranged from under 4 GPa to just over 5 GPa. Most species had an average elastic modulus between 90 GPa and 110 GPa, with three specimens with values below and two above this range (Table 1). TED and MC ranged from 2.3 g/cm^3^ to 2.6 g/cm^3^, and 2 g/cm^3^ to 2.4 g/cm^3^, respectively. Average prism width ranged from 3.5 μm (rhesus macaque upper molar) to 5.7 μm (gorilla lower molar). Values for each variable (except prism size, see below) varied across the five crown positions, with common patterns observed among samples. Mean values for each of the five positions are presented for each sample in Supplementary Tables 2-4, by variable type. Many variables were significantly associated with others, including elastic modulus with TED/MC and hardness, and TED/HC with prism width (Figure 2; Supplementary Table 5). There was no significant association between enamel thickness and any parameters investigated (Supplementary Table 5). Similarly, there was no increase in enamel hardness with increase in enamel thickness. Body mass was also not associated with any of the variables studied, with the exception of mean prism width, in which larger prism size was associated with larger body mass (r(18) = .64, p = .004).

**Table 1.**
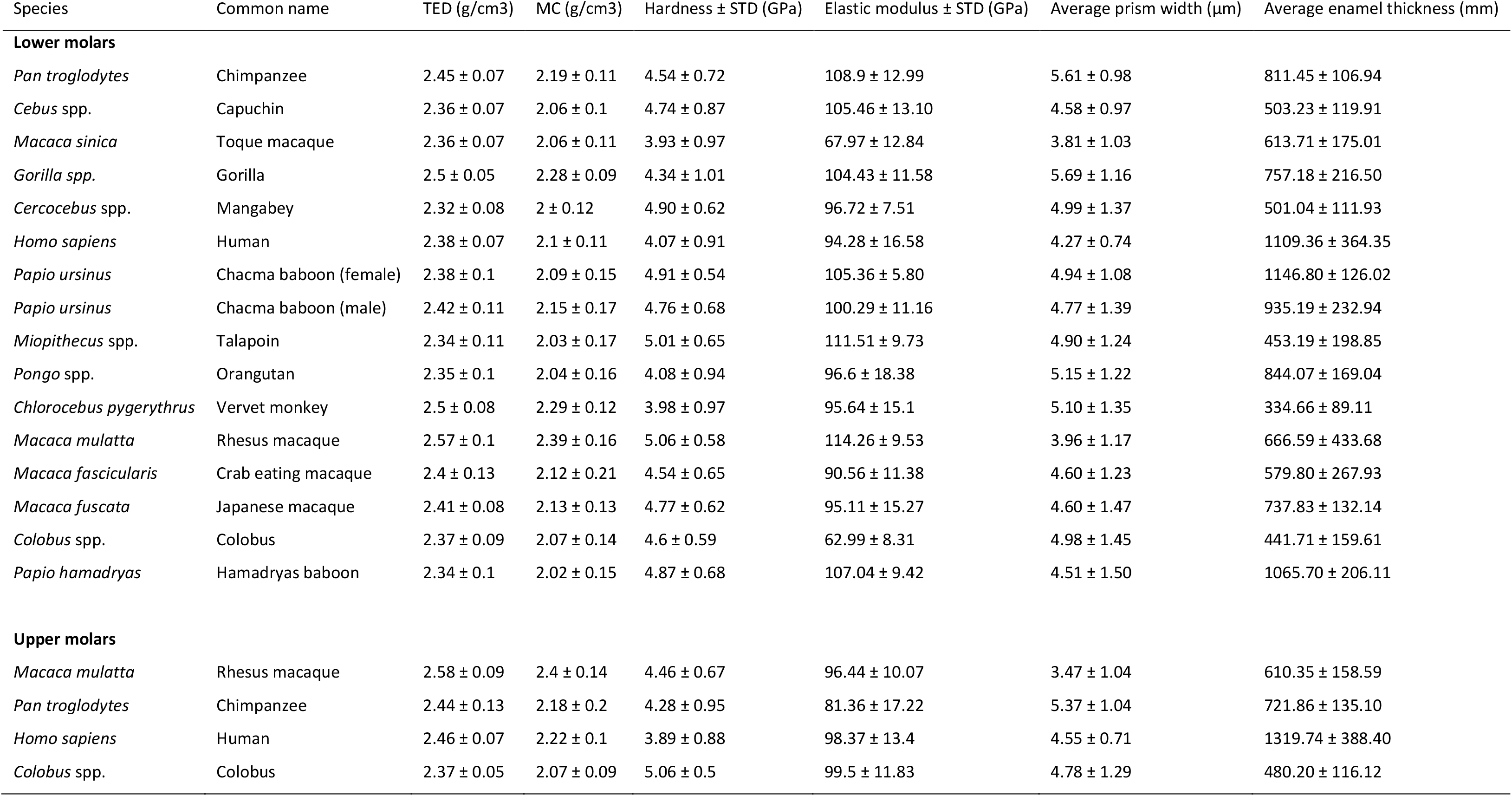
Species averages for each variable studied. TED: total effective density; MC: mineral concentration.

| Species | Common name | TED (g/cm3) | MC (g/cm3) | Hardness ± STD (GPa) | Elastic modulus ± STD (GPa) | Average prism width (μm) | Average enamel thickness (mm) |
| --- | --- | --- | --- | --- | --- | --- | --- |
| **Lower molars** |  |  |  |  |  |  |  |
| *Pan troglodytes* | Chimpanzee | 2.45 ± 0.07 | 2.19 ± 0.11 | 4.54 ± 0.72 | 108.9 ± 12.99 | 5.61 ± 0.98 | 811.45 ± 106.94 |
| *Cebus* spp. | Capuchin | 2.36 ± 0.07 | 2.06 ± 0.1 | 4.74 ± 0.87 | 105.46 ± 13.10 | 4.58 ± 0.97 | 503.23 ± 119.91 |
| *Macaca sinica* | Toque macaque | 2.36 ± 0.07 | 2.06 ± 0.11 | 3.93 ± 0.97 | 67.97 ± 12.84 | 3.81 ± 1.03 | 613.71 ± 175.01 |
| *Gorilla* spp. | Gorilla | 2.5 ± 0.05 | 2.28 ± 0.09 | 4.34 ± 1.01 | 104.43 ± 11.58 | 5.69 ± 1.16 | 757.18 ± 216.50 |
| *Cercocebus* spp. | Mangabey | 2.32 ± 0.08 | 2 ± 0.12 | 4.90 ± 0.62 | 96.72 ± 7.51 | 4.99 ± 1.37 | 501.04 ± 111.93 |
| *Homo sapiens* | Human | 2.38 ± 0.07 | 2.1 ± 0.11 | 4.07 ± 0.91 | 94.28 ± 16.58 | 4.27 ± 0.74 | 1109.36 ± 364.35 |
| *Papio ursinus* | Chacma baboon (female) | 2.38 ± 0.1 | 2.09 ± 0.15 | 4.91 ± 0.54 | 105.36 ± 5.80 | 4.94 ± 1.08 | 1146.80 ± 126.02 |
| *Papio ursinus* | Chacma baboon (male) | 2.42 ± 0.11 | 2.15 ± 0.17 | 4.76 ± 0.68 | 100.29 ± 11.16 | 4.77 ± 1.39 | 935.19 ± 232.94 |
| *Miopithecus* spp. | Talapoin | 2.34 ± 0.11 | 2.03 ± 0.17 | 5.01 ± 0.65 | 111.51 ± 9.73 | 4.90 ± 1.24 | 453.19 ± 198.85 |
| *Pongo* spp. | Orangutan | 2.35 ± 0.1 | 2.04 ± 0.16 | 4.08 ± 0.94 | 96.6 ± 18.38 | 5.15 ± 1.22 | 844.07 ± 169.04 |
| *Chlorocebus pygerythrus* | Vervet monkey | 2.5 ± 0.08 | 2.29 ± 0.12 | 3.98 ± 0.97 | 95.64 ± 15.1 | 5.10 ± 1.35 | 334.66 ± 89.11 |
| *Macaca mulatta* | Rhesus macaque | 2.57 ± 0.1 | 2.39 ± 0.16 | 5.06 ± 0.58 | 114.26 ± 9.53 | 3.96 ± 1.17 | 666.59 ± 433.68 |
| *Macaca fascicularis* | Crab eating macaque | 2.4 ± 0.13 | 2.12 ± 0.21 | 4.54 ± 0.65 | 90.56 ± 11.38 | 4.60 ± 1.23 | 579.80 ± 267.93 |
| *Macaca fuscata* | Japanese macaque | 2.41 ± 0.08 | 2.13 ± 0.13 | 4.77 ± 0.62 | 95.11 ± 15.27 | 4.60 ± 1.47 | 737.83 ± 132.14 |
| *Colobus* spp. | Colobus | 2.37 ± 0.09 | 2.07 ± 0.14 | 4.6 ± 0.59 | 62.99 ± 8.31 | 4.98 ± 1.45 | 441.71 ± 159.61 |
| *Papio hamadryas* | Hamadryas baboon | 2.34 ± 0.1 | 2.02 ± 0.15 | 4.87 ± 0.68 | 107.04 ± 9.42 | 4.51 ± 1.50 | 1065.70 ± 206.11 |
| **Upper molars** |  |  |  |  |  |  |  |
| *Macaca mulatta* | Rhesus macaque | 2.58 ± 0.09 | 2.4 ± 0.14 | 4.46 ± 0.67 | 96.44 ± 10.07 | 3.47 ± 1.04 | 610.35 ± 158.59 |
| *Pan troglodytes* | Chimpanzee | 2.44 ± 0.13 | 2.18 ± 0.2 | 4.28 ± 0.95 | 81.36 ± 17.22 | 5.37 ± 1.04 | 721.86 ± 135.10 |
| *Homo sapiens* | Human | 2.46 ± 0.07 | 2.22 ± 0.1 | 3.89 ± 0.88 | 98.37 ± 13.4 | 4.55 ± 0.71 | 1319.74 ± 388.40 |
| *Colobus* spp. | Colobus | 2.37 ± 0.05 | 2.07 ± 0.09 | 5.06 ± 0.5 | 99.5 ± 11.83 | 4.78 ± 1.29 | 480.20 ± 116.12 |

**Figure 2.**
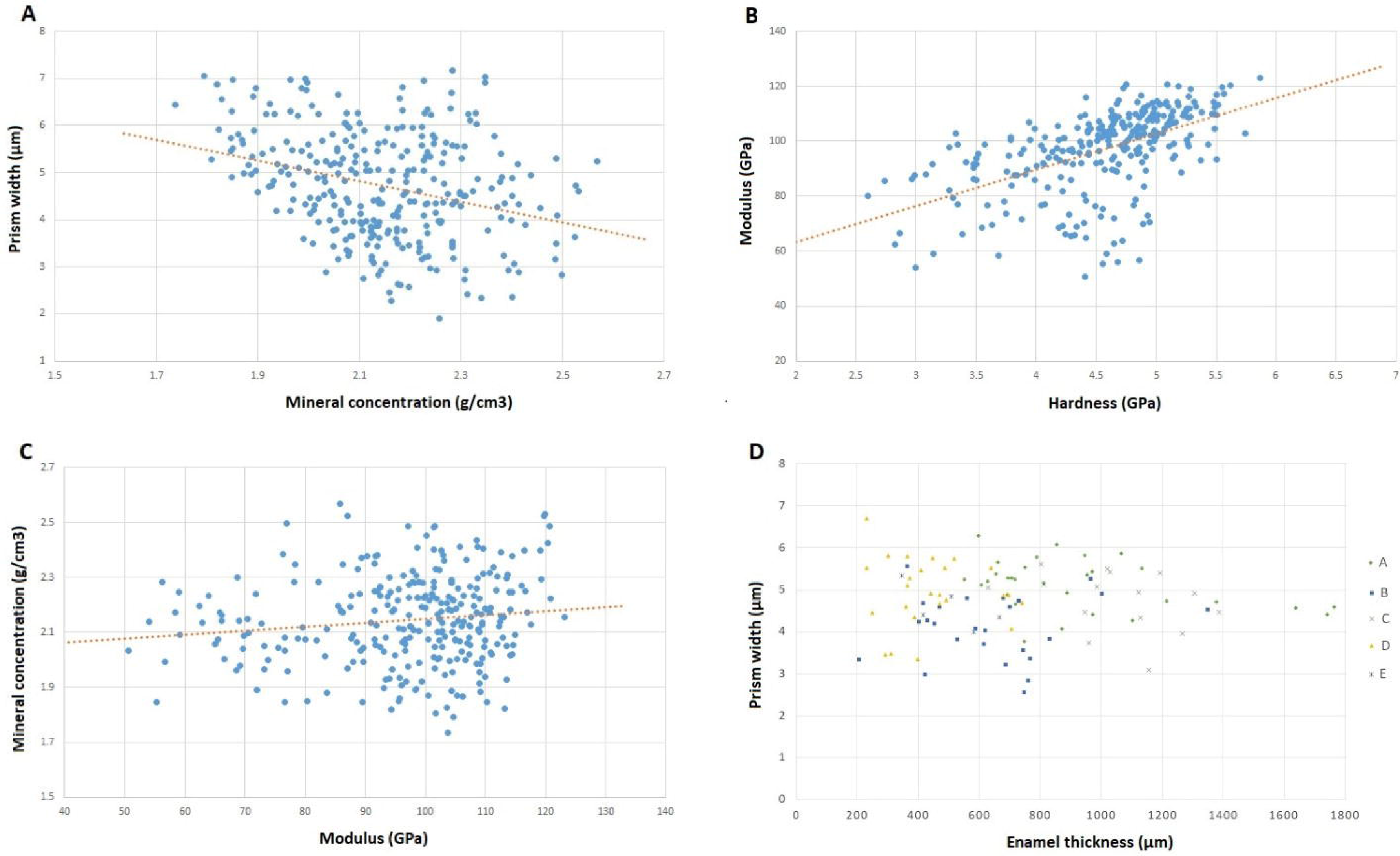
Plots comparing variables studied, including A) prism size and mineral concentration; B) elastic modulus vs hardness; C) mineral concentration vs elastic modulus; D) average prism width vs enamel thickness, for each of the five positions in each specimen studied, split into the following groups: apes (A), macaques (B); baboons (C), all other Catarrhini (D), and Capuchin (E).

Non-functional sides of teeth (buccal positions in upper molars and lingual in lower molars) had higher mechanical properties values than their functional counterparts (lingual positions in upper molars and buccal in lower molars) in most comparisons between the same positions on opposite sides of the tooth (Figures 3 and 4). When the overall data was compiled for functional vs. non-functional sides for both upper and lower teeth, there were significantly higher values for hardness for non-functional positions, with a mean of 4.5 GPa ± 0.67 for functional and 4.71 GPa ± 0.53 for non-functional sides (t(110) = −2.57, p = .011). When only lower molars were compared for hardness values, there was still a significant difference for functional vs. non-functional sides (t(86) = 2.54, p = .012). The mean hardness values for buccal lateral enamel hardness was 4.47 GPa ± 0.73, and for lingual lateral enamel in lower molars it was higher with a mean of 4.69 ± 0.46, although this difference between sides was not significant (t(42) = −1.669, p = .099). In lower molars, hardness values for buccal cuspal enamel were significantly lower than for lingual cuspal enamel (buccal cuspal: 4.60 GPa ± 0.64; lingual cuspal: 4.82 GPa ± 0.39; (t(44) = −1.994, p = .049).

**Figure 3.**
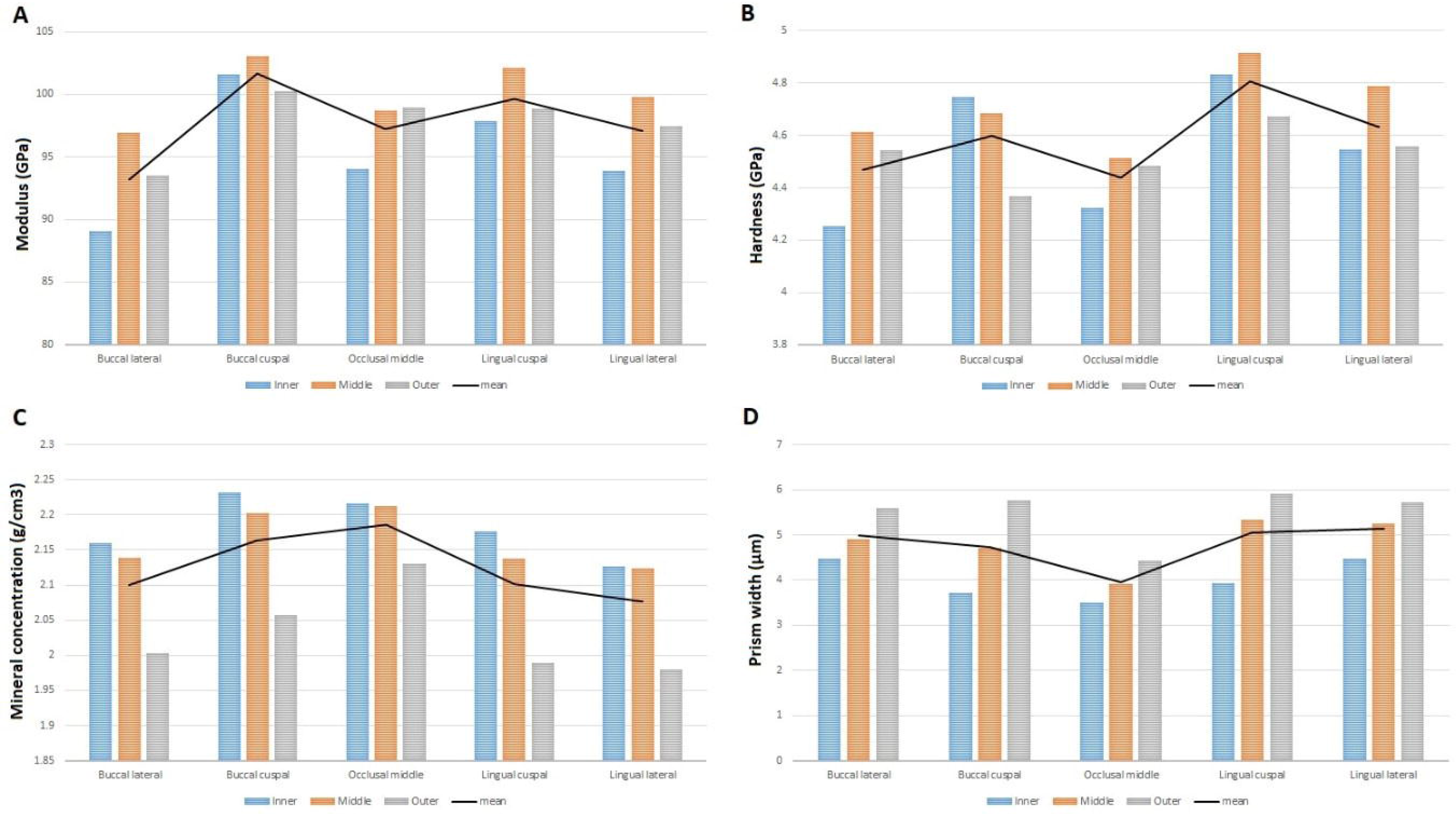
Lower molar comparisons. Mean inner, middle and outer values for each of the five positions studied, showing an overall mean trend line for A) elastic modulus (GPa); B) hardness (GPa); C) mineral concentration (g/cm^3^); D) prism width (μm).

**Figure 4.**
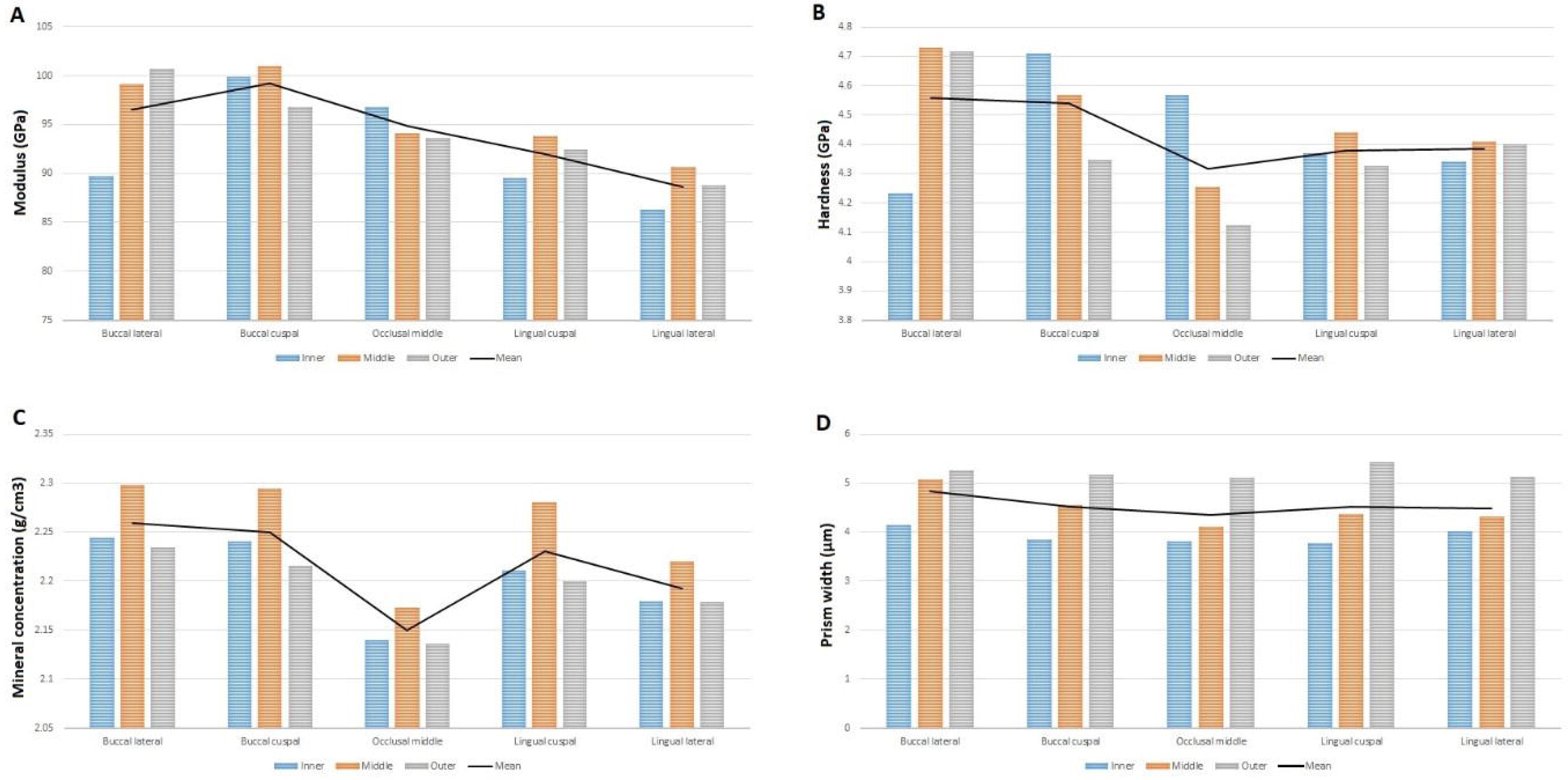
Upper molar comparisons. Mean inner, middle and outer values for each of the five positions studied, showing an overall mean trend line for A) elastic modulus (GPa); B) hardness (GPa); C) mineral concentration (g/cm^3^); D) prism width (μm).

For elastic modulus values, there was no significant differences between functional vs. non-functional sides (t(110) = −1.632, p = .104). Although functional sides had lower average values (functional sides: 95.97 GPa ± 17.05; non-functional: 99.20 GPa ± 11.92). Similarly, there were no statistical differences in elastic modulus when only lower molars were compared (t(86) = 0.885, p = .377) or when spilt by specific positions, for example lateral positions for lower teeth (t(41) = −1.341, p = . 184), and the cuspal positions for lower teeth (t(44) = 0.146, p = .885).

Prism size increased from inner to outer enamel in all samples studied (Supplementary Table 6). When inner, middle and outer averages for each specimen were compared, inner enamel prism width was significantly smaller than outer enamel (t(19) = −7.342, p < .001). Inner prism width was also significantly smaller than in middle enamel (t(19) = −3.929, p < .001), and likewise middle prism width was smaller than outer enamel (t(19) = −3.537, p = .001). When the five positions were compared in different species, there was no significant relationship between enamel thickness and prism size (Figure 2D; r(100) = −.026, p = .805). Average prism size was relatively constant across the five different crown positions, with the exception of occlusal middle enamel (Figure 3D; Figure 4D). Typically, the inner enamel within the cusp regions (buccal cuspal and lingual cuspal) was characterised by small prisms, while the outer enamel had larger prisms (Figure 3D and 4D). Lateral enamel was characterised by a lower range of prism sizes from inner to outer enamel, with a larger prism size in inner enamel than in cuspal positions. The occlusal middle enamel had smaller prism sizes in each of the three categories (inner, middle and outer).

Middle enamel was most commonly the hardest (Table 2). When all samples were compiled, the middle enamel had significantly higher hardness values than the outer enamel (t(95) = 2.113, p = .036), with no other significant differences found between the three positions. Although the elastic modulus was also higher in the middle position (Table 2), this was only significantly higher when compared to inner enamel (t(95) = −2.037, p = .043). Mechanical property values in cuspal enamel were typically higher than in lateral enamel, with nine of the twelve comparisons for elastic modulus, hardness and TED/MC showing higher values in cusps than in lateral enamel (Supplementary Tables 3 and 4).

**Table 2.** Average elastic modulus (GPa) and hardness (GPa) for each specimen, split by inner, middle and outer enamel positions.

| | Modulus ± STD (GPa) | | | Hardness ± STD (GPa) | | |
| --- | --- | --- | --- | --- | --- | --- |
|  | Inner | Middle | Outer | Inner | Middle | Outer |
| **Lower molars** |  |  |  |  |  |  |
| Chimpanzee | 104.02 ± 14.09 | 112.15 ± 13.03 | 110.39 ± 10.76 | 4.55 ± 0.78 | 4.58 ± 0.78 | 4.48 ± 0.64 |
| Capuchin | 107.23 ± 15.93 | 105.81 ± 13.74 | 103.25 ± 13.07 | 4.85 ± 1 | 4.81 ± 0.91 | 4.57 ± 0.87 |
| Toque macaque | 70.53 ± 12.74 | 74.51 ± 10.28 | 71.99 ± 11.25 | 4 ± 0.96 | 4.34 ± 0.78 | 4.22 ± 0.84 |
| Gorilla | 102.72 ± 11.67 | 108.39 ± 10.86 | 102.28 ± 11.66 | 4.15 ± 1.08 | 4.58 ± 1.03 | 4.29 ± 0.83 |
| Mangabey | 94.57 ± 8.57 | 98.51 ± 6.79 | 97.49 ± 6.76 | 4.83 ± 0.77 | 5.02 ± 0.54 | 4.83 ± 0.46 |
| Human | 98.33 ± 14.38 | 100.22 ± 15.66 | 85.27 ± 15.9 | 4.13 ± 1.19 | 4.28 ± 0.74 | 3.88 ± 0.8 |
| Chacma baboon (female) | 106.51 ± 5.88 | 105.9 ± 5.5 | 103.24 ± 5.58 | 5.12 ± 0.43 | 4.95 ± 0.52 | 4.59 ± 0.56 |
| Chacma baboon (male) | 100.52 ± 11.49 | 101.77 ± 10.65 | 98.13 ± 10.93 | 4.94 ± 0.66 | 4.79 ± 0.64 | 4.49 ± 0.66 |
| Talapoin | 112.61 ± 9.31 | 112.86 ± 6.97 | 109.03 ± 11.24 | 5.15 ± 0.57 | 5.04 ± 0.64 | 4.84 ± 0.68 |
| Orangutan | 84.69 ± 20.67 | 100.59 ± 15.25 | 104.89 ± 12.31 | 3.49 ± 1.02 | 4.31 ± 0.9 | 4.46 ± 0.63 |
| Vervet monkey | 93.48 ± 11.39 | 97.21 ± 13.89 | 95.56 ± 18.01 | 3.93 ± 0.99 | 4.06 ± 1 | 3.91 ± 0.93 |
| Rhesus macaque | 113.57 ± 8.78 | 114.19 ± 10.65 | 114.79 ± 8.7 | 5.24 ± 0.54 | 4.98 ± 0.65 | 4.97 ± 0.5 |
| Crab eating macaque | 80.64 ± 9 | 95.32 ± 8.07 | 96.17 ± 8.81 | 4.02 ± 0.56 | 4.88 ± 0.53 | 4.75 ± 0.43 |
| Japanese macaque | 90.85 ± 15.48 | 97.47 ± 12.97 | 94.79 ± 17.93 | 4.74 ± 0.76 | 4.83 ± 0.49 | 4.66 ± 0.62 |
| Colobus | 58.95 ± 7.66 | 65.88 ± 6.74 | 66.39 ± 8.3 | 4.54 ± 0.5 | 4.78 ± 0.46 | 4.54 ± 0.67 |
| Hamadryas baboon | 105.17 ± 12.26 | 108.95 ± 8.4 | 106.43 ± 5.49 | 4.97 ± 0.88 | 4.96 ± 0.58 | 4.69 ± 0.48 |
| **Upper molars** |  |  |  |  |  |  |
| Rhesus macaque | 96.38 ± 12.67 | 97.4 ± 9.28 | 94.43 ± 8.78 | 4.55 ± 0.78 | 4.52 ± 0.61 | 4.28 ± 0.6 |
| Chimpanzee | 78.08 ± 17.54 | 80.73 ± 18.53 | 87.7 ± 15.92 | 4.21 ± 1.06 | 4.13 ± 1.05 | 4.62 ± 0.74 |
| Human | 99.69 ± 12.92 | 101.33 ± 13.33 | 95.62 ± 14.9 | 4.03 ± 0.99 | 4 ± 0.88 | 3.72 ± 0.79 |
| Colobus | 96.18 ± 14.74 | 103.06 ± 9.18 | 100.61 ± 9.89 | 5.05 ± 0.54 | 5.24 ± 0.41 | 4.93 ± 0.49 |

All samples studied here displayed a *Schmelzmuster* with Hunter-Schreger Bands in all positions, with the exception of occlusal middle enamel in which visualization of HSB was sometimes difficult. HSB were typically present in the inner ¾ of the enamel thickness (Figure 5). In some teeth these bands extended further, covering almost the entire length of the enamel thickness (Figure 5a). Radial enamel made up the approximate ¼ of the outer enamel in many samples (Figure 5b). Therefore, most inner and middle indentations were taken within Hunter-Schreger bands, whereas outer readings would have often been indented on radial enamel. Species averages for hardness, elastic modulus, and TED/MC all fell within a relatively narrow range of values. Based on the small sample sizes for individual groups, detailed phylogenetic comparisons were difficult.

**Figure 5.**
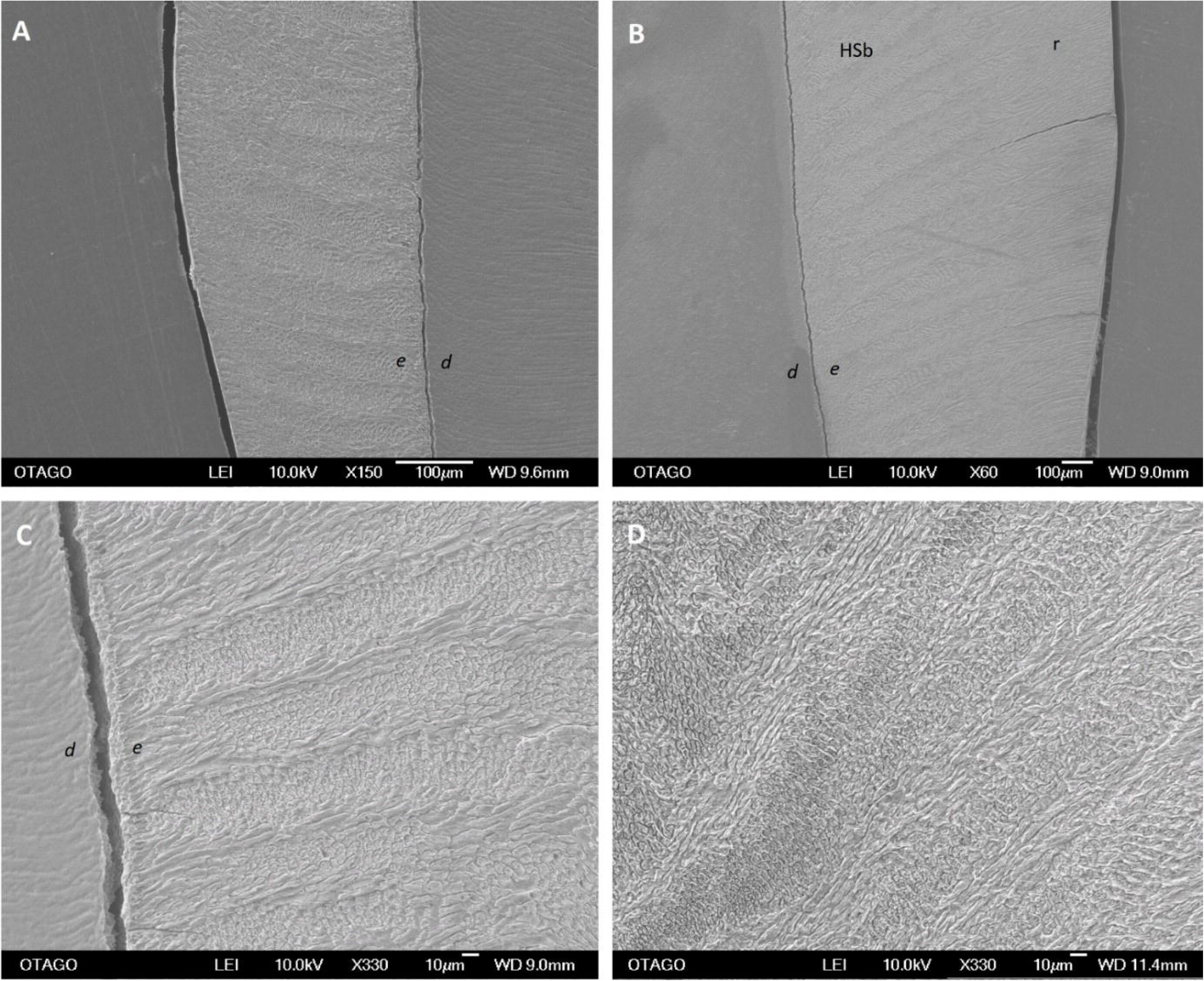
Enamel structure in different species, showing the presence of Hunter-Schreger bands in lower molars A) vervet monkey, lingual lateral position, showing Hunter-Schreger bands stretching almost the entire enamel thickness; B) hamadryas baboon, lingual lateral position, showing Hunter-Schreger bands extending approximately half to two thirds the way across the enamel thickness C) colobus, buccal lateral, close up of Hunter-Schreger bands in the inner enamel; D) talapoin, lingual cuspal, showing Hunter-Schreger bands. d=dentine; e=enamel; HSb=Hunter-Schreger bands; r=radial enamel.

## Discussion

Middle enamel had the highest hardness and elastic modulus values, and non-functional molar sides were significantly harder than their functional counterparts. Prism size and mineral density values showed a significant relationship with mechanical properties in certain comparisons, but variation in enamel structures should also be accounted. From a biomechanical perspective, primate enamel is not homogeneous, both across the extent of the enamel thickness (i.e., inner to outer enamel) as well as among specific locations of the tooth (e.g., buccal Vs. lingual and cuspal Vs., lateral).

This study had some limitations. Most studies on the mechanical properties of teeth have relied on dry specimens (e.g., Baker et al., 1959; Sanson et al., 2007; Braly et al., 2007; Lee et al., 2010; Constantino et al., 2012; Erickson, 2014). Other studies have used ‘fresh’ (also called ‘wet’ or ‘native’) specimens, to better represent the real mechanical properties of enamel during life, with differences in values reported between these two methods (e.g., Kaiser et al., 2018). Here we studied dry museum specimens, and therefore the mechanical property readings may be slightly higher than the true-life values. However, since we are interested in variations across tooth crowns, and the comparative literature also used dry samples, this would not have an effect on the results and conclusions reported here, since all specimens were preserved in a similar manner.

Mechanical properties of enamel may also vary depending on the scale and orientation in which the tooth is studied, due to the anisotropic nature of enamel. Since the positions were standardized in the present study, this likely reduced these effects. Additionally, due to substantial proportions of the enamel thickness being composed of Hunter-Schreger Bands in all samples studied, the prism orientation in relation to the nanoindenter tip would have fluctuated regularly. Differences in elastic modulus and hardness have also been reported as higher in enamel prism areas than in interprismatic regions (Ang et al., 2012), therefore likely also influencing results. In this study, we did not attempt to assess the orientation of the prisms or the coverage of prismatic/interprismatic enamel that was indented. Instead, we explored overall position averages. More detailed studies may allow further inferences on the underlying causes for the patterns described here.

The first hypothesis that hardness and elastic modulus values will increase with increase enamel thickness was not supported since there was no association between enamel thickness and hardness and elastic modulus values. Similarly, enamel density variables (TED and MC) were also not associated with enamel thickness. There seems to be no evidence to suggest functional sides of teeth had adaptations to ‘strengthen’ this side of the tooth. In fact, the opposite was observed for hardness values, with higher hardness values in non-functional cusps. There seems to be no support for the second hypothesis, which stated that mechanical property and density values would increase from inner to outer enamel. In this study, mid-enamel values were higher for both elastic modulus and hardness. There was however an increase in average prism size from inner to outer enamel, and the largest differences were observed in areas of thicker enamel, with the two cusp positions (lingual and buccal cuspal) showing the largest mean differences between inner and outer positions. The third hypothesis that cuspal enamel will also show higher values for enamel properties than lateral enamel was supported for all variables, except for prism size.

Although variation in mechanical properties across primate crowns was observed in the present study, this was within a narrow range of values. This variation was much less than that described in Cuy et al. (2002), in which the hardness of human enamel at the surface averaged 4.6 GPa, down to 3.4 GPa near the EDJ. In the current study, the differences in hardness values between outer and inner enamel were under 0.5 GPa. This was also supported by the Campbell et al. (2012) study on lemurs and humans, which included similar methods to those used here (section through buccal-lingual in mesial cusps). In humans, substantial differences in mechanical properties between inner and outer enamel was observed both in hardness and elastic modulus values, but the same was not true for other primates, with either no trend observed or a more minor variation. Constantino et al. (2012), also observed this difference between clinical human samples and dental samples from extant primates. They found some primates did not show the steep increase in mechanical property values from inner to outer enamel. They concluded that “in several species, the point of maximum modulus or hardness appears not at either the inner or outer surface but somewhere in between” (Constantino et al., 2012, page 174). The present study confirms this association, and goes further by suggesting this may in fact be the most common pattern found in primates, at least in Catarrhini.

Why human molars (at least for clinical samples) display such significant variation in mechanical property values, increasing from inner to outer enamel, requires further investigation. Mechanical properties of enamel may change during the life of an individual, and environmental and dietary factors may also influence enamel mechanical properties. Park et al (2008b) observed an increase in both hardness and elastic modulus near the occlusal surface in older humans, although the inner enamel remained relatively constant among age groups. It is possible that studies that observed higher mechanical property values in outer enamel may relate at least partly to the age differences of sampled individuals. Most of the teeth analyzed here were from young individuals, since in wild primates only these individuals have relatively complete molars (i.e., before significant tooth wear occurs). This may help explain differences in enamel properties between humans and other primates.

Prism width is larger in non-functional positions (the two upper buccal positions and two lower lingual positions). The reasons for this need to be explored further, however this might be related to differences in mechanical properties between these positions. Although mineral concentration and prism size likely affect the mechanical properties of enamel, other structural and compositional properties may also influence property differences observed across primate tooth crowns. The results of this study support research on human enamel which has shown outer enamel lacks the microstructural features of inner/middle enamel which helps dissipate fracture energy (Bajaj and Arola, 2009). However, why inner/middle enamel, that has more interprismatic enamel, has higher hardness readings in most primates requires further investigation. A possible explanation is the presence of larger scale enamel structures such as Hunter-Schreger Bands, which are found in all species studied and typically located from inner to middle enamel. There are however a variety of other factors that likely influence the observed variation in mechanical properties, including prism orientation, enamel composition, and crystal arrangement/orientation (e.g., Cuy et al., 2002; Angker et al., 2004; Shimizu and Macho, 2008; Xie et al., 2009; Jeng et al., 2011; Stifler et al., 2021).

The results of this study suggest that variation in mechanical properties values across molar crowns may relate to responses to localized stresses and specific functions. Previous studies have shown that non-functional cusps had higher prevalence of fractures, but a low rate of wear, compared to their functional counterparts (Cavel et al., 1985; Eakle et al., 1986; Towle et al., 2021; Towle and Loch, 2021). Based on the results of the present study, non-functional cusps had higher hardness values, as a potential response to avoiding fracture. The thickly enameled functional cusps would need more material to stay functional for longer periods (i.e., as the tooth gradually wears down). Why the middle enamel had higher elastic modulus and hardness values needs to be explored further, but there are a variety of studies that have highlighted the potential benefits of evolving graded enamel, and this likely relates to protecting teeth against different types of failure (e.g., He and Swain, 2009; He et al., 2013).

Species averages for hardness, elastic modulus, and TED/MC all fall within a relatively narrow range of values. Based on the small sample for individual groups, detailed phylogenetic comparisons are difficult, however some interesting results justify future investigation. For example, when buccal and lingual positions are combined for lower molars, only five teeth show higher hardness values on the buccal side (out of 16). Of these five, all three baboon samples are included. A similar pattern was observed for inner, middle and outer enamel, with only six of the 16 teeth having the highest mean hardness values for inner enamel. Again, all three baboon specimens were included in these six teeth. Lastly, in lower molars only three teeth show the highest values for hardness in the buccal lateral location, and all these were baboons.

Similarly, apes had low levels of hardness but moderate to high elastic modulus values. For example, out of the five ape teeth studied, the highest hardness value was 4.54 GPa, yet in the three baboon teeth the lowest value was 4.76 GPa. Similar to earlier studies, among apes, humans had the lowest hardness values, followed by gorillas and chimpanzees. Therefore, despite relatively low variation in mechanical properties among the species studied, there are still minor differences observable among groups. The relationship between mechanical properties of enamel and dietary variation in primates remains opaque. There does not seem to be differences in mechanical properties among frugivore or folivore species, and groups that have often been considered hard-object feeders (e.g., capuchin, mangabey and orangutan) do not show elevated hardness and elastic modulus values compared to others. As it has been suggested previously (e.g., Lee et al., 2010; Constantino et al., 2012), it seems other enamel properties have evolved in response to changes in food processing behaviours (e.g., occlusal morphology and enamel thickness). This is supported by the fact that enamel thickness was not associated with prism width, but body mass was. This likely relates to allometric factors, and prism size seems to be conserved. In contrast, enamel thickness can change rapidly in response to dietary changes (Thiery et al., 2017).

This study adds evidence for variation in mechanical properties in primate crowns, further highlighting that enamel is not a homogeneous material. Non-functional molar sides may have evolved to be stronger due to additional localised forces. As well as prism size and mineral concentration, the composition and structure variation (e.g., Hunter-Schreger Bands), may also be crucial in explaining variation in mechanical properties. The little variation in mechanical properties values and mineral concertation among species may suggest these characteristics are evolutionary conserved, and likely shared among Catarrhini and primates more generally. We did not find evidence for rapid change in mechanical properties of enamel associated with dietary differences within closely related primate groups, suggesting other features (e.g., enamel thickness and occlusal morphology) had a greater influence in recent adaptation. However, slight but discernable variations in properties among taxa may allow more precise biomechanical and phylogenetic interpretations and inferences for the groups studied.

## Acknowledgements

This research was supported by a Sir Thomas Kay Sidey Postdoctoral fellowship from the Faculty of Dentistry University of Otago to Ian Towle. We thank E. Girvan and A. McNaughton from Otago OMNI for assistance during SEM and micro-CT analyses. The authors thank the Study Material Committee from the Primate Research Institute (PRI), Kyoto University, for access to their collections, and T. Ito for assistance during data collection. The research was partly performed under the Cooperative Research Program of the PRI (2019-C-20). We also thank the museums in New Zealand that provided samples for this study and the curators for all their assistance, including Otago Museum (E. Burns), Museum of Natural Mystery (B. Mahalski), and Auckland Museum (M. Rayner and R. Moore).

---

**Supplementary Table 1.** Samples studied, including species, tooth type, sex, specimen number and collection.

| Species | Common name | Tooth | Collection | Specimen number | Sex |
| --- | --- | --- | --- | --- | --- |
| *Papio ursinus* | Chacma baboon | Lower right M2 | Museum of Natural Mystery | n/a | Male |
| *Papio ursinus* | Chacma baboon | Lower right M2 | Museum of Natural Mystery | n/a | Female |
| *Chlorocebus pygerythrus* | Vervet monkey | Lower left M2 | Museum of Natural Mystery | n/a | Male |
| *Macaca fascicularis* | Crab eating macaque | Lower right M1 | Museum of Natural Mystery | n/a | Female |
| *Macaca fuscata* | Japanese macaque | Lower left M2 | Primate Research Institute | PRI 3383 | Male |
| *Papio hamadryas* | Hamadryas baboon | Lower right M2 | Primate Research Institute | PRI 7148 | Male |
| *Pan troglodytes* | Chimpanzee | Lower right M1 | Auckland Museum | LM 315 | Not known |
| *Pan troglodytes* | Chimpanzee | Upper left M1 | Auckland Museum | LM 315 | Not known |
| *Colobus* spp. | Colobus | Lower left M2 | Auckland Museum | LM 957 | Not known |
| *Colobus* spp. | Colobus | Upper left M2 | Auckland Museum | LM 957 | Not known |
| *Macaca mulatta* | Rhesus macaque | Lower left M2 | Dental Faculty | CU 138 | Female |
| *Macaca mulatta* | Rhesus macaque | Upper left M2 | Dental Faculty | CU 138 | Female |
| *Homo sapiens* | Human | Lower left M2 | Dental Faculty | n/a | Not known |
| *Homo sapiens* | Human | Upper left M2 | Dental Faculty | n/a | Not known |
| *Miopithecus* spp. | Talapoin | Lower right M2 | Otago Museum | VT 2702 | Not known |
| *Gorilla* spp. | Gorilla | Lower right M1 | Otago Museum | VT 2468 | Male |
| *Pongo* spp. | Orangutan | Lower right M1 | Otago Museum | VT 3361 | Not known |
| *Cercocebus* spp. | Mangabey | Lower right M1 | Otago Museum | VT 2715 | Not known |
| *Macaca sinica* | Toque macaque | Lower right M2 | Otago Museum | VT 2707 | Not known |
| *Cebus* spp. | Capuchin | Lower left M2 | Otago Museum | VT 2728 | Not known |

**Supplementary Table 2.** Enamel properties data. Mean prism width (left value) and enamel thickness (right value) for each position studied. All measurements are in μm.

|  | Species | Buccal lateral | Buccal cuspal | Occlusal middle | Lingual cuspal | Lingual lateral |
| --- | --- | --- | --- | --- | --- | --- |
| **Lower molars** | Chimpanzee | 6.08 ± 1.06; 854.52 ± 106.57 | 5.36 ± 1.12; 954.67 ± 34.31 | 5.29 ± 0.78; 707.21 ± 80.05 | 5.79 ± 1.01; 789.6 ± 36.43 | 5.54 ± 0.68; 751.27 ± 22.75 |
| | Capuchin | 4.39 ± 0.80; 415.92 ± 43.05 | 4.34 ± 0.67; 664.67 ± 60.31 | 3.99 ± 0.66; 581.46 ± 17.18 | 4.84 ± 1.05; 508.44 ± 9.59 | 5.34 ± 1.05; 345.65 ± 23.60 |
| | Toque macaque | 4.18 ± 0.78; 453.19 ± 54.45 | 3.82 ± 1.19; 832.39 ± 58.93 | 2.83 ± 0.63; 761.68 ± 56.43 | 4.02 ± 0.70; 618.97 ± 13.63 | 4.22 ± 1.11; 402.3 ± 34.16 |
| | Gorilla | 5.24 ± 1.53; 550.55 ± 33.53 | 5.88 ± 1.13; 1066.25 ± 11.85 | 5.21 ± 0.83; 626.87 ± 112.79 | 5.82 ± 1.03; 945.78 ± 15.72 | 6.29 ± 0.83; 596.45 ± 17.27 |
| | Mangabey | 5.53 ± 0.69; 486.46 ± 41.51 | 4.87 ± 1.32; 695.36 ± 12.36 | 3.34 ± 0.66; 397.7 ± 27.85 | 5.75 ± 1.58; 516.56 ± 12.47 | 5.47 ± 0.75; 409.13 ± 41.20 |
| | Human | 3.77 ± 0.56; 748.45 ± 91.00 | 4.40 ± 0.67; 1740.61 ± 37.97 | 4.06 ± 0.73; 871.66 ± 135.05 | 4.73 ± 0.84; 1214.02 ± 68.26 | 4.41 ± 0.51; 972.07 ± 82.33 |
| | Chacma baboon (female) | 5.08 ± 1.52; 985.37 ± 53.83 | 4.92 ± 0.89; 1305.05 ± 59.62 | 4.33 ± 0.51; 1128.66 ± 128.65 | 4.94 ± 0.87; 1122.11 ± 42.95 | 5.41 ± 1.15; 1192.8 ± 38.02 |
| | Chacma baboon (male) | 5.05 ± 1.15; 628.16 ± 108.31 | 3.95 ± 0.89; 1266.11 ± 65.63 | 3.73 ± 0.63; 959.51 ± 58.76 | 5.51 ± 1.60; 1019.58 ± 72.87 | 5.61 ± 1.26; 802.57 ± 126.88 |
| | Talapoin | 5.53 ± 0.85; 230.95 ± 3.07 | 5.52 ± 1.56; 638.31 ± 46.24 | 4.06 ± 0.56; 705.73 ± 54.79 | 4.92 ± 1.44; 440.68 ± 16.92 | 4.46 ± 0.79; 250.3 ± 10.37 |
| | Orangutan | 5.16 ± 0.71; 812.71 ± 47.03 | 5.52 ± 1.63; 1132.15 ± 14.76 | 4.89 ± 0.94; 692.12 ± 48.77 | 5.29 ± 1.37; 694.38 ± 27.34 | 4.92 ± 1.18; 889 ± 62.57 |
| | Vervet monkey | 6.71 ± 0.96; 231.38 ± 22.85 | 4.89 ± 0.82; 469.6 ± 86.46 | 3.48 ± 0.57; 310.99 ± 19.29 | 4.60 ± 0.59; 360.22 ± 5.16 | 5.81 ± 0.95; 301.1 ± 12.14 |
| | Rhesus macaque | 3.33 ± 0.82; 206.73 ± 15.99 | 4.52 ± 0.89; 1349.8 ± 448.61 | 2.56 ± 0.50; 747.12 ± 57.51 | 4.80 ± 0.96; 560.12 ± 32.01 | 4.59 ± 0.75; 469.16 ± 37.69 |
| | Crab eating macaque | 5.56 ± 1.28; 364.01 ± 37.06 | 4.90 ± 1.19; 1003.14 ± 316.76 | 3.80 ± 0.91; 528.18 ± 52.14 | 4.06 ± 1.21; 586.99 ± 40.72 | 4.67 ± 0.67; 416.69 ± 36.19 |
| | Japanese macaque | 4.73 ± 1.18; 729.68 ± 35.71 | 3.70 ± 1.29; 614.34 ± 31.75 | 5.26 ± 1.86; 965.87 ± 109.37 | 4.79 ± 1.40; 678.74 ± 16.61 | 4.58 ± 1.21; 700.51 ± 17.94 |
| | Colobus | 5.10 ± 0.87; 363.92 ± 20.59 | 4.68 ± 1.70; 739.71 ± 33.55 | 3.45 ± 0.82; 293 ± 6.16 | 5.76 ± 1.36; 446.99 ± 2.67 | 5.81 ± 0.90; 364.91 ± 23.49 |
| | Hamadryas baboon | 4.46 ± 1.02; 946.88 ± 53.19 | 4.46 ± 1.03; 1386.26 ± 51.80 | 3.09 ± 0.58; 1155.4 ± 111.67 | 5.43 ± 1.68; 1028.45 ± 11.38 | 5.13 ± 1.72; 811.49 ± 32.07 |
| **Upper molars** | Rhesus macaque | 4.27 ± 1.25; 429.44 ± 22.22 | 3.55 ± 0.81; 744.95 ± 15.89 | 2.97 ± 1.14; 422.53 ± 6.42 | 3.21 ± 0.88; 687.38 ± 37.57 | 3.36 ± 0.54; 767.46 ± 70.81 |
| | Chimpanzee | 5.11 ± 0.90; 605.14 ± 21.61 | 5.25 ± 0.85; 717.88 ± 56.36 | 5.38 ± 0.63; 654.68 ± 26.36 | 5.43 ± 1.12; 970.58 ± 31.90 | 5.66 ± 1.49; 661.03 ± 31.57 |
| | Human | 4.65 ± 0.66; 719.06 ± 53.17 | 4.56 ± 0.54; 1637.94 ± 31.59 | 4.70 ± 0.68; 1376.46 ± 84.65 | 4.58 ± 0.68; 1763.64 ± 137.08 | 4.27 ± 0.88; 1101.62 ± 52.46 |
| | Colobus | 5.28 ± 0.92; 372.58 ± 14.35 | 4.75 ± 1.34; 491.49 ± 7.70 | 4.34 ± 1.42; 387.94 ± 9.17 | 4.87 ± 1.66; 679.38 ± 55.83 | 4.66 ± 0.78; 469.63 ± 57.26 |

**Supplementary Table 3.** Mechanical properties data. Mean elastic modulus (left value) and hardness (right value) for each position studied. All measurements are in GPa.

| Upper/lower | Species | Buccal lateral | Buccal cuspal | Occlusal middle | Lingual cuspal | Lingual lateral |
| --- | --- | --- | --- | --- | --- | --- |
| **Lower molars** | Chimpanzee | - | 105.14 ± 6.93; 4.38 ± 0.57 | 114.70 ± 13.52; 4.71 ± 0.68 | 111.05 ± 16.13; 4.57 ± 0.70 | 105.25 ± 13.37; 4.56 ± 0.94 |
|  | Capuchin | 93.2 ± 14.68; 4.07 ± 0.94 | 117.22 ± 11.37; 5.54 ± 0.67 | 108.34 ± 5.49; 4.69 ± 0.6 | 108.89 ± 9.12; 4.81 ± 0.66 | 97.77 ± 7.84; 4.46 ± 0.75 |
|  | Toque macaque | 58.4 ± 13.77; 3.5 ± 1.03 | 62.46 ± 4.85; 2.32 ± 0.35 | 68.55 ± 6.44; 4.07 ± 0.69 | 77.5 ± 7.73; 4.45 ± 0.74 | 72.93 ± 12.57; 3.91 ± 1.07 |
|  | Gorilla | 106.74 ± 15.67; 4.54 ± 1.32 | 102.99 ± 12.97; 3.53 ± 0.9 | - | 101.39 ± 6.71; 4.66 ± 0.41 | 106.22 ± 7.21; 4.71 ± 0.6 |
|  | Mangabey | 95.23 ± 8.7; 4.52 ± 0.62 | 101.13 ± 5.14; 4.63 ± 0.7 | 97.15 ± 7.26; 5.3 ± 0.5 | 92.89 ± 7.59; 4.84 ± 0.6 | 97.88 ± 6.47; 4.97 ± 0.44 |
|  | Human | 85.67 ± 29.34; 3.46 ± 0.87 | 98.23 ± 15.02; 4.45 ± 0.73 | 92.42 ± 8.71; 3.82 ± 0.84 | 95.43 ± 12.67; 4.11 ± 0.84 | 97.34 ± 13.28; 4.39 ± 1.02 |
|  | Chacma baboon (female) | 107.08 ± 3.62; 5.14 ± 0.27 | 104.96 ± 5.23; 4.86 ± 0.58 | 106.06 ± 7.6; 4.88 ± 0.58 | 106.27 ± 5.03; 5.09 ± 0.54 | 102.44 ± 5.58; 4.59 ± 0.34 |
|  | Chacma baboon (male) | 112.54 ± 4.89; 5.16 ± 0.35 | 106.74 ± 4.78; 4.83 ± 0.58 | 90.97 ± 11.3; 4.18 ± 0.71 | 94.8 ± 5.97; 4.84 ± 0.47 | 96.08 ± 8.38; 4.84 ± 0.82 |
|  | Talapoin | 112.61 ± 8.08; 5.18 ± 0.68 | 112.93 ± 8.52; 5 ± 0.65 | 108.76 ± 8.05; 4.74 ± 0.49 | 112.03 ± 12.33; 4.96 ± 0.77 | 111.45 ± 10.7; 5.26 ± 0.52 |
|  | Orangutan | 79.99 ± 17.59; 3.42 ± 0.82 | 99.06 ± 15.38; 3.9 ± 0.86 | 108.18 ± 12.89; 4.46 ± 0.81 | 98.34 ± 18.31; 4.63 ± 0.7 | 89.08 ± 18.88; 3.98 ± 1.06 |
|  | Vervet monkey | 85.7 ± 5.86; 3.5 ± 0.41 | 90.95 ± 5.42; 3.45 ± 0.29 | 86.65 ± 7.05; 2.91 ± 0.44 | 112.32 ± 19.94; 5.11 ± 0.67 | 98.28 ± 8.27; 4.75 ± 0.66 |
|  | Rhesus macaque | 110.13 ± 8.66; 5.12 ± 0.6 | 120.7 ± 7.74; 5.29 ± 0.34 | 113.24 ± 10.05; 4.7 ± 0.61 | 117.66 ± 6.09; 5.27 ± 0.43 | 108.75 ± 10.23; 4.97 ± 0.65 |
|  | Crab eating macaque | 84.49 ± 11.03; 4.34 ± 0.63 | 86.18 ± 9.5; 4.71 ± 0.37 | 99.41 ± 8.57; 4.57 ± 0.54 | 91.65 ± 12.36; 4.59 ± 0.9 | 90.95 ± 9.18; 4.53 ± 0.64 |
|  | Japanese macaque | 69.2 ± 7.77; 4.37 ± 0.57 | 106 ± 7.63; 4.88 ± 0.54 | 98.9 ± 9.88; 4.84 ± 0.49 | 99.1 ± 11.91; 4.96 ± 0.66 | 96.66 ± 8.17; 4.65 ± 0.67 |
|  | Colobus | 53.98 ± 4.5; 4.59 ± 0.48 | 61.54 ± 5.34; 4.59 ± 0.37 | 70.62 ± 4.12; 4.54 ± 0.54 | 69.39 ± 8.36; 4.92 ± 0.8 | 65.39 ± 7.74; 4.56 ± 0.8 |
|  | Hamadryas baboon | 110.09 ± 5.48; 5.34 ± 0.36 | 110.47 ± 7.72; 4.87 ± 0.61 | 97.25 ± 13.05; 4.09 ± 0.67 | 107.91 ± 4.96; 5.07 ± 0.47 | 108.6 ± 7.2; 4.98 ± 0.58 |
| **Upper molars** | Rhesus macaque | 83.11 ± 14.95; 3.83 ± 0.81 | 102.52 ± 4.94; 4.72 ± 0.52 | 95.64 ± 7.45; 4.39 ± 0.57 | 99.7 ± 6.42; 4.68 ± 0.68 | 96.56 ± 7.59; 4.42 ± 0.54 |
|  | Chimpanzee | 103.5 ± 8.13; 4.95 ± 0.61 | 94.99 ± 6.58; 5.07 ± 0.49 | 80.32 ± 6.58; 4.47 ± 0.67 | 61.47 ± 9.41; 3.36 ± 0.74 | 70.97 ± 8.43; 3.77 ± 0.85 |
|  | Human | 98.96 ± 12.53; 3.89 ± 0.83 | 91.04 ± 10.38; 3.23 ± 0.55 | 99.4 ± 18.6; 3.55 ± 0.76 | 102.18 ± 10.2; 4.45 ± 0.81 | 101.5 ± 12.17; 4.36 ± 0.74 |
|  | Colobus | 100.13 ± 6.23; 5.54 ± 0.3 | 106.93 ± 10.2; 5.06 ± 0.54 | 104.49 ± 5.22; 4.89 ± 0.42 | 102.74 ± 8.21; 4.91 ± 0.39 | 83.72 ± 11.06; 4.92 ± 0.53 |

**Supplementary Table 4.** Mineral density data. Mean mineral density (left value) and total effective density (right value) for each position studied. All measurements are in g/cm^3^.

| Upper/lower | Species | Buccal lateral | Buccal cuspal | Occlusal middle | Lingual cuspal | Lingual lateral |
| --- | --- | --- | --- | --- | --- | --- |
| **Lower molars** | Chimpanzee | 2.12 ± 0.13; 2.4 ± 0.08 | 2.2 ± 0.08; 2.45 ± 0.05 | 2.29 ± 0.07; 2.5 ± 0.05 | 2.19 ± 0.07; 2.44 ± 0.05 | 2.18 ± 0.09; 2.44 ± 0.06 |
|  | Capuchin | 2.02 ± 0.08; 2.34 ± 0.05 | 2.1 ± 0.1; 2.39 ± 0.06 | 2.1 ± 0.09; 2.38 ± 0.06 | 2.02 ± 0.12; 2.33 ± 0.08 | 2.05 ± 0.09; 2.35 ± 0.06 |
|  | Toque macaque | 2.06 ± 0.11; 2.36 ± 0.07 | 2.05 ± 0.14; 2.35 ± 0.09 | 2.14 ± 0.1; 2.41 ± 0.06 | 2.03 ± 0.11; 2.34 ± 0.07 | 2.02 ± 0.07; 2.34 ± 0.04 |
|  | Gorilla | 2.25 ± 0.07; 2.48 ± 0.04 | 2.32 ± 0.1; 2.53 ± 0.07 | 2.28 ± 0.09; 2.5 ± 0.06 | 2.3 ± 0.06; 2.51 ± 0.04 | 2.27 ± 0.08; 2.49 ± 0.05 |
|  | Mangabey | 1.99 ± 0.1; 2.31 ± 0.06 | 2.02 ± 0.11; 2.33 ± 0.07 | 2.06 ± 0.14; 2.36 ± 0.09 | 1.95 ± 0.13; 2.29 ± 0.08 | 1.97 ± 0.11; 2.3 ± 0.07 |
|  | Human | 2.07 ± 0.06; 2.37 ± 0.04 | 2.16 ± 0.07; 2.42 ± 0.04 | 2.19 ± 0.06; 2.44 ± 0.04 | 2.1 ± 0.07; 2.38 ± 0.05 | 1.96 ± 0.07; 2.3 ± 0.04 |
|  | Chacma baboon (female) | 2.13 ± 0.1; 2.41 ± 0.06 | 2.02 ± 0.12; 2.33 ± 0.08 | 2.25 ± 0.11; 2.48 ± 0.07 | 1.96 ± 0.13; 2.3 ± 0.08 | 2.11 ± 0.1; 2.39 ± 0.07 |
|  | Chacma baboon (male) | 2.23 ± 0.05; 2.47 ± 0.03 | 2.1 ± 0.15; 2.38 ± 0.09 | 2.33 ± 0.1; 2.53 ± 0.06 | 1.95 ± 0.11; 2.29 ± 0.07 | 2.14 ± 0.12; 2.41 ± 0.07 |
|  | Talapoin | 1.9 ± 0.22; 2.26 ± 0.14 | 2.1 ± 0.12; 2.39 ± 0.08 | 2.17 ± 0.11; 2.43 ± 0.07 | 2.02 ± 0.11; 2.33 ± 0.07 | 1.97 ± 0.12; 2.31 ± 0.08 |
|  | Orangutan | 2.07 ± 0.13; 2.37 ± 0.08 | 2.03 ± 0.11; 2.34 ± 0.07 | 2.17 ± 0.14; 2.43 ± 0.09 | 2.03 ± 0.15; 2.34 ± 0.09 | 1.89 ± 0.14; 2.25 ± 0.09 |
|  | Vervet monkey | 2.32 ± 0.06; 2.52 ± 0.04 | 2.34 ± 0.11; 2.54 ± 0.07 | 2.2 ± 0.11; 2.45 ± 0.07 | 2.33 ± 0.15; 2.53 ± 0.1 | 2.23 ± 0.11; 2.47 ± 0.07 |
|  | Rhesus macaque | 2.35 ± 0.16; 2.55 ± 0.1 | 2.51 ± 0.18; 2.65 ± 0.12 | 2.3 ± 0.11; 2.51 ± 0.07 | 2.36 ± 0.18; 2.55 ± 0.12 | 2.4 ± 0.08; 2.58 ± 0.05 |
|  | Crab eating macaque | 1.95 ± 0.12; 2.29 ± 0.08 | 2.31 ± 0.15; 2.52 ± 0.1 | 2.08 ± 0.12; 2.37 ± 0.08 | 2.25 ± 0.18; 2.48 ± 0.11 | 2 ± 0.19; 2.32 ± 0.12 |
|  | Japanese macaque | 2.09 ± 0.1; 2.38 ± 0.06 | 2.16 ± 0.12; 2.42 ± 0.08 | 2.21 ± 0.07; 2.46 ± 0.04 | 2.17 ± 0.11; 2.43 ± 0.07 | 2.03 ± 0.12; 2.34 ± 0.08 |
|  | Colobus | 1.96 ± 0.13; 2.3 ± 0.08 | 2.19 ± 0.14; 2.44 ± 0.09 | 2.11 ± 0.08; 2.39 ± 0.05 | 2.04 ± 0.13; 2.35 ± 0.08 | 2.05 ± 0.09; 2.35 ± 0.06 |
|  | Hamadryas baboon | 2.09 ± 0.12; 2.38 ± 0.07 | 2.01 ± 0.14; 2.33 ± 0.09 | 2.1 ± 0.14; 2.39 ± 0.09 | 1.94 ± 0.16; 2.29 ± 0.1 | 1.97 ± 0.15; 2.3 ± 0.09 |
| **Upper molars** | Rhesus macaque | 2.53 ± 0.11; 2.66 ± 0.07 | 2.45 ± 0.11; 2.61 ± 0.07 | 2.29 ± 0.09; 2.51 ± 0.06 | 2.46 ± 0.12; 2.61 ± 0.07 | 2.27 ± 0.06; 2.5 ± 0.04 |
|  | Chimpanzee | 2.32 ± 0.1; 2.52 ± 0.06 | 2.24 ± 0.09; 2.48 ± 0.06 | 1.86 ± 0.16; 2.23 ± 0.1 | 2.21 ± 0.11; 2.45 ± 0.07 | 2.26 ± 0.13; 2.49 ± 0.08 |
|  | Human | 2.18 ± 0.06; 2.43 ± 0.04 | 2.19 ± 0.1; 2.44 ± 0.06 | 2.34 ± 0.1; 2.54 ± 0.06 | 2.18 ± 0.09; 2.43 ± 0.06 | 2.19 ± 0.05; 2.44 ± 0.03 |
|  | Colobus | 2.01 ± 0.08; 2.33 ± 0.05 | 2.12 ± 0.05; 2.4 ± 0.03 | 2.11 ± 0.08; 2.39 ± 0.05 | 2.08 ± 0.09; 2.37 ± 0.06 | 2.04 ± 0.08; 2.35 ± 0.05 |

**Supplementary Table 5.** Results of statistical test of relationship between the different variables studied, using a Pearson correlation coefficient. The individual data points are the individual mean readings for inner, middle and outer for the 5 positions in the 20 teeth. The exception was comparisons with enamel thickness which use mean values for each position. Bold figures indicate significance.

|  | Hardness | Elastic modulus | Ted/MC | Average Prism width |
| --- | --- | --- | --- | --- |
| Hardness | - | **r(288) = .5421, p < .00001** | r(288) = -.0731, p = .216789 | r(288) = -.0061, p = .919248 |
| Elastic modulus | **r(288) = .5421, p < .00001** | - | **r(288) = .1313, p = .025867** | r(288) = .0203, p = .731569 |
| Ted/MC | r(288) = -.0731, p = .216789 | **r(288) = .1313, p = .025867** | - | **r(300) = -0.3072, p < .00001** |
| Average Prism width | r(288) = -.0061, p = .919248 | r(288) = .0203, p = .731569 | **r(300) = -0.3072, p < .00001** | - |
| Average enamel thickness SEM | r(98) = -0.1392, p = .17225 | r(98) = 0.1687, p = .096806 | r(100) = 0.0798, p = .429979 | r(100) = -.0259, p = .804989 |

**Supplementary Table 6.** Prism width (μm), average total effective density (g/cm^3^) and average mineral concentration (g/cm^3^), split by inner, middle and outer enamel for each specimen.

| | Prism Width ($\mu\text{m}$) | | | Average Total Effective Density ($\text{g}/\text{cm}^3$) | | | Average Mineral Concentration ($\text{g}/\text{cm}^3$) | | |
| --- | --- | --- | --- | --- | --- | --- | --- | --- | --- |
|  | Inner | Middle | Outer | Inner | Middle | Outer | Inner | Middle | Outer |
| **Lower molars** |  |  |  |  |  |  |  |  |  |
| Chimpanzee | 4.90 ± 0.79 | 5.83 ± 0.92 | 6.10 ± 0.80 | 2.48 ± 0.03 | 2.47 ± 0.04 | 2.39 ± 0.08 | 2.25 ± 0.05 | 2.23 ± 0.07 | 2.1 ± 0.12 |
| Capuchin | 3.89 ± 0.69 | 4.63 ± 0.82 | 5.21 ± 0.91 | 2.4 ± 0.04 | 2.38 ± 0.04 | 2.29 ± 0.06 | 2.12 ± 0.06 | 2.1 ± 0.06 | 1.95 ± 0.1 |
| Toque macaque | 3.035 ± 0.67 | 3.89 ± 1.06 | 4.52 ± 0.74 | 2.4 ± 0.05 | 2.38 ± 0.04 | 2.3 ± 0.07 | 2.12 ± 0.09 | 2.09 ± 0.07 | 1.96 ± 0.11 |
| Gorilla | 5.06 ± 1.09 | 5.67 ± 0.98 | 6.33 ± 1.07 | 2.51 ± 0.04 | 2.53 ± 0.04 | 2.46 ± 0.05 | 2.3 ± 0.06 | 2.32 ± 0.06 | 2.21 ± 0.08 |
| Mangabey | 4.56 ± 0.94 | 4.66 ± 1.23 | 5.75 ± 1.57 | 2.35 ± 0.06 | 2.34 ± 0.06 | 2.26 ± 0.07 | 2.05 ± 0.1 | 2.03 ± 0.09 | 1.9 ± 0.12 |
| Human | 3.9 ± 0.60 | 4.25 ± 0.75 | 4.67 ± 0.67 | 2.39 ± 0.06 | 2.4 ± 0.06 | 2.36 ± 0.07 | 2.1 ± 0.09 | 2.13 ± 0.09 | 2.06 ± 0.12 |
| Chacma baboon (female) | 4.00 ± 0.56 | 5.34 ± 0.83 | 5.48 ± 1.10 | 2.43 ± 0.05 | 2.4 ± 0.08 | 2.31 ± 0.09 | 2.18 ± 0.08 | 2.13 ± 0.12 | 1.98 ± 0.15 |
| Chacma baboon (male) | 3.77 ± 0.91 | 4.66 ± 1.05 | 5.88 ± 1.27 | 2.47 ± 0.07 | 2.43 ± 0.09 | 2.35 ± 0.11 | 2.23 ± 0.12 | 2.18 ± 0.14 | 2.04 ± 0.18 |
| Talapoin | 4.06 ± 0.84 | 4.97 ± 1.06 | 5.66 ± 1.25 | 2.37 ± 0.1 | 2.37 ± 0.1 | 2.29 ± 0.11 | 2.07 ± 0.16 | 2.07 ± 0.15 | 1.95 ± 0.18 |
| Orangutan | 4.05 ± 0.75 | 5.38 ± 0.98 | 6.04 ± 0.93 | 2.42 ± 0.06 | 2.37 ± 0.07 | 2.26 ± 0.09 | 2.15 ± 0.1 | 2.08 ± 0.11 | 1.9 ± 0.15 |
| Vervet monkey | 4.66 ± 1.10 | 5.28 ± 1.50 | 5.34 ± 1.35 | 2.52 ± 0.08 | 2.53 ± 0.07 | 2.46 ± 0.07 | 2.32 ± 0.12 | 2.33 ± 0.11 | 2.21 ± 0.11 |
| Rhesus macaque | 3.40 ± 0.87 | 3.94 ± 1.22 | 4.53 ± 1.14 | 2.61 ± 0.08 | 2.59 ± 0.09 | 2.53 ± 0.12 | 2.45 ± 0.13 | 2.42 ± 0.15 | 2.32 ± 0.18 |
| Crab eating macaque | 3.93 ± 1.04 | 4.74 ± 1.40 | 5.10 ± 0.92 | 2.44 ± 0.12 | 2.42 ± 0.12 | 2.34 ± 0.14 | 2.18 ± 0.19 | 2.16 ± 0.19 | 2.03 ± 0.22 |
| Japanese macaque | 3.28 ± 0.59 | 4.88 ± 1.51 | 5.59 ± 1.04 | 2.45 ± 0.04 | 2.43 ± 0.05 | 2.34 ± 0.09 | 2.2 ± 0.07 | 2.17 ± 0.08 | 2.03 ± 0.14 |
| Colobus | 4.29 ± 1.18 | 4.73 ± 1.39 | 5.92 ± 1.27 | 2.41 ± 0.07 | 2.39 ± 0.07 | 2.3 ± 0.09 | 2.14 ± 0.11 | 2.11 ± 0.11 | 1.96 ± 0.14 |
| Hamadryas baboon | 3.53 ± 0.76 | 4.34 ± 1.23 | 5.68 ± 1.54 | 2.39 ± 0.05 | 2.36 ± 0.06 | 2.25 ± 0.1 | 2.1 ± 0.08 | 2.07 ± 0.1 | 1.89 ± 0.17 |
| **Upper molars** |  |  |  |  |  |  |  |  |  |
| Rhesus macaque | 2.58 ± 0.63 | 3.61 ± 0.97 | 4.21 ± 0.76 | 2.56 ± 0.09 | 2.59 ± 0.09 | 2.58 ± 0.09 | 2.38 ± 0.14 | 2.43 ± 0.15 | 2.4 ± 0.14 |
| Chimpanzee | 4.62 ± 0.55 | 5.35 ± 0.73 | 6.14 ± 1.13 | 2.42 ± 0.1 | 2.45 ± 0.14 | 2.45 ± 0.13 | 2.16 ± 0.16 | 2.21 ± 0.21 | 2.2 ± 0.21 |
| Human | 4.17 ± 0.71 | 4.65 ± 0.63 | 4.83 ± 0.62 | 2.45 ± 0.04 | 2.5 ± 0.06 | 2.42 ± 0.07 | 2.2 ± 0.07 | 2.28 ± 0.09 | 2.16 ± 0.11 |
| Colobus | 4.29 ± 0.81 | 4.35 ± 1.26 | 5.70 ± 1.22 | 2.38 ± 0.05 | 2.39 ± 0.04 | 2.33 ± 0.06 | 2.1 ± 0.07 | 2.1 ± 0.07 | 2.01 ± 0.09 |

